# Midfrontal theta and pupil dilation parametrically track subjective conflict (but also surprise) during intertemporal choice

**DOI:** 10.1101/172122

**Authors:** Hause Lin, Blair Saunders, Cendri A Hutcherson, Michael Inzlicht

**Author notes:** Corresponding author Hause Lin Department of Psychology, University of Toronto 1265 Military Trail Toronto, ON, M1C 1A4, Canada.

## Abstract

Many everyday choices are based on personal, subjective preferences. When choosing between two options, we often feel conflicted, especially when trading off costs and benefits occurring at different times (e.g., saving for later versus spending now). Although previous work has investigated the neurophysiological basis of conflict during inhibitory control tasks, less is known about subjective conflict resulting from competing subjective preferences. In this pre-registered study, we investigated subjective conflict during intertemporal choice, whereby participants chose between smaller immediate versus larger delayed rewards (e.g., $15 today vs. $22 in 30 days). We used economic modeling to parametrically vary eleven different levels of conflict, and recorded EEG data and pupil dilation. Midfrontal theta power, derived from EEG, correlated with pupil responses, and our results suggest that these signals track different gradations of subjective conflict. Unexpectedly, both signals were also maximally enhanced when decisions were surprisingly easy. Therefore, these signals may track events requiring increased attention and adaptive shifts in behavioral responses, with subjective conflict being only one type of such event. Our results suggest that the neural systems underlying midfrontal theta and pupil responses interact when weighing costs and benefits during intertemporal choice. Thus, understanding these interactions might elucidate how individuals resolve self-control conflicts.

**Highlights:**

- Modeled conflict during intertemporal choice and measured EEG and pupil responses.
- Midfrontal theta and pupil responses parametrically tracked subjective conflict.
- But theta and pupil responses were also large when decisions were surprisingly easy.
- These signals may implement adaptive control during value-guided choice.

## 1. Introduction

Many everyday decisions are value-guided. People have preferences—they value choice options to varying degrees—and then decide based on these subjective preferences (e.g., I prefer Android over iPhones, savory over sweet foods, or spending now over saving for later). At the neural level, the brain assigns a subjective value to each available choice option and compares these values to arrive at a choice (Camerer, 2013; Kable and Glimcher, 2009; Konovalov and Krajbich, 2016; Rangel et al., 2008). Because of the cost-benefit trade-offs involved in choosing between options, decision makers often feel conflicted when making value-guided decisions (Frank et al., 2015; Shenhav and Buckner, 2014).

### 1.1. Objective versus subjective conflict

What we currently know about the neural correlates of decision conflict has largely been informed by studies using inhibitory control tasks (e.g., Stroop, go/no-go), whereby decisions are determined by objective states of the world (e.g., Botvinick et al., 2001). For example, on an incompatible (high-conflict) Stroop trial, reading the printed word would be incorrect but reading the color in which the word is printed in would be correct. Recent work, however, suggests that decisions based on objective states and subjective preferences involve slightly different processes (Polanía et al., 2014; Pisauro et al., 2017; Summerfield and Tsetsos, 2012). Thus, drawing inferences about how people arbitrate between two closely valued options (e.g., Android vs. iPhone) from inhibitory control studies (e.g., is that word presented in red vs. green font) might be premature. Here, we examine the neurophysiological correlates of conflict resulting from competing subjective preferences using the classic intertemporal choice (delay discounting) task (Ainslie, 1975; Frederick et al., 2002; Thaler, 1981).

The intertemporal choice paradigm has been extensively used to investigate the psychological and neural underpinnings of self-control dilemmas and subjective value representation (Bernhardt et al., 2014; McClure et al., 2004; Zauberman and Urminsky, 2016). Many economic models accurately describe intertemporal preferences, which have been associated with real-life behaviors requiring self-control, including pathological gambling, substance abuse, and social media usage (Dixon et al, 2003; Kollins, 2003; Shenhav et al., 2017). Crucially, this paradigm has often been used to investigate how different neural systems may contribute to competing valuations that give rise to self-control conflicts (Berns et al., 2007; McClure et al., 2007). Here, we investigate how different gradations of subjective conflict during intertemporal choice parametrically modulate two neurophysiological signals, namely midfrontal theta power and pupil dilation.

### 1.2. Midfrontal EEG conflict signals

Electroencephalography (EEG) is a temporally precise technique that is often used to investigate how conflict-related neural activity evolves over time (Cohen, 2017; Frank et al., 2005; Yeung et al., 2004). Previous work suggests that conflict-related activity originates from the dorsal anterior cingulate cortex (dACC) and surrounding medial prefrontal cortical (mPFC) regions (Debener et al., 2005; Töllner et al., 2017; Ebitz and Platt, 2015). Consistent with these findings, theta oscillations (~4−8 Hz) measured over midfrontal EEG electrode sites increase during high conflict trials (Cavanagh et al., 2012; Frank et al., 2015), and these theta dynamics are thought to implement adaptive control processes necessary for resolving conflict (Cavanagh and Frank, 2014; Cohen, 2014b; Verguts, 2017). In addition, theta oscillations might underlie event-related components (ERPs) such as the N2 and error-related negativity that are observed when conflicting, mutually exclusive responses are activated simultaneously (e.g., Cavanagh et al., 2012; Yeung et al., 2004).

EEG studies of conflict often focus on objective response conflict (e.g., Botvinick et al., 2001; Töllner et al., 2017; Yeung et al., 2004), and the studies that have investigated subjective conflict have typically used reinforcement learning paradigms whereby participants have to initially learn to associate novel cues with different reward probabilities and subsequently (during a test phase) try to accurately choose cues that have been previously associated with higher reward probabilities (e.g., Cavanagh et al., 2011; Frank et al., 2015; but see also Nakao et al., 2010; Pisauro et al., 2017). Here, we focus on subjective conflict during an intertemporal choice task, which only requires participants to express their pre-existing preferences for rewards presented at different delays. That is, little or no learning is required during the task, and participants will only be indicating their natural preferences, which cannot be classified as accurate or inaccurate. Thus, our task will allow us to investigate whether and how EEG signals measured over midfrontal scalp electrodes track subjective conflict even when there lack clear, objectively accurate answers.

### 1.3. Conflict-related pupil dilation responses

Brain regions commonly implicated in value-guided choice include the orbitofrontal cortex and ACC, and these regions interconnect strongly with the brainstem nucleus locus coeruleus, which releases norepinephrine to modulate neural gain to optimize decision making (Aston-Jones and Cohen, 2005a; Aston-Jones and Cohen, 2005b; Aston-Jones and Waterhouse, 2016; Berridge and Waterhouse, 2003; Eldar et al., 2013). Although locus coeruleus-norepinephrine (LC-NE) gain-adjustment activity has been proposed to interact with EEG signals (Cavanagh and Frank, 2014; Nieuwenhuis et al., 2005; Nieuwenhuis et al., 2011; Singer 2013; Verguts and Notebaert, 2009; Womelsdorf et al., 2014), it remains a challenge to study these proposed interactions because locus coeruleus activity can be difficult to measure in humans.

Pupil diameter, however, appears to be a promising noninvasive correlate of locus coeruleus activity and neural gain (Aston-Jones and Cohen, 2005b; Joshi et al., 2016; Murphy et al., 2011; Murphy et al., 2014; Rajkowski et al., 1994). For example, changes in pupil size have been associated with behaviors, such as exploit-explore trade-offs, that have been associated with LC-NE system activity and gain adjustment (Eldar et al., 2013; Eldar et al., 2016; Gilzenrat et al., 2010; Jepma and Nieuwenhuis, 2011; Murphy et al., 2016; Warren et al., 2016). In addition, pupil responses are ideal for investigating conflict-related processes because they correlate with increased attention or mental effort during decision making (Kahneman & Beatty, 1966; Siegle et al., 2003; Simpson, 1969), autonomic arousal and ACC activity during inhibitory control task performance (Critchley et al., 2005; Laeng et al., 2011), and conflict during decision making (Cavanagh et al., 2014). Critically, recent studies have provided indirect evidence for interactions between LC-NE and EEG signals by showing that pupil dilation correlates with EEG signals (e.g., theta oscillations) during perceptual and inhibitory control tasks (Dippel et al., 2017; Hong et al., 2014; Mückschel et al., 2016; Mückschel et al., 2017).

However, whether theta-pupil relationships are also present during value-guided choice remains untested. We therefore investigated how changes in pupil responses relate to different levels of subjective conflict, and whether pupil responses correlate with midfrontal theta power during intertemporal choice. Showing such consilience—that similar basic processes are conserved across different types of decisions—will indicate not only that theta-pupil associations generalize across choice domains, but also that these correlations are robust and replicable, an issue of renewed importance in psychology and neuroscience (Button et al., 2013; Cohen, 2017; Crandall and Sherman, 2016; Open Science Collaboration, 2015; Smaldino and McElreath, 2016).

### 1.4. Present Study

Using a pre-registered modeling approach and parametric design (osf.io/7m9c2), we investigated whether subjective conflict during intertemporal choice parametrically modulates midfrontal theta and pupil responses. Our study also extends recent work showing theta-pupil correlations during inhibitory control tasks (Dippel et al., 2017).

Participants made intertemporal decisions several days before the main experiment. We then fitted the hyperbolic discounting model to each participant's data (Green and Myerson, 2004), and parametrically varied intertemporal preferences and subjective choice conflict separately for each participant during a subsequent laboratory session (i.e., by using each participant's discount function to generate participant-specific delayed rewards). Previous work used this neurometric approach to identify brain regions that encode the subjective value of delayed rewards during intertemporal choice (Kable and Glimcher, 2007; Peters and Büchel, 2010). Here, we further show that by modeling intertemporal preferences, we can provide insights into the neurophysiology of subjective conflict and self-control dilemmas involving cost-benefit trade-offs.

During the main task, we concurrently recorded EEG activity and pupil dilation as participants performed an intertemporal choice task with individually-tailored and model-derived delayed rewards. Our results show that although theta power measured over midfrontal scalp electrodes and pupil dilation responses seem to parametrically track subjective conflict during intertemporal choice, both signals were, unexpectedly, enhanced when the decisions were surprisingly easy and involved little or no conflict. Thus, conflict itself may not be required to evoke midfrontal theta and pupil dilation responses. Our findings are consistent with past work suggesting that midfrontal theta is evoked by events that require increased attention and adaptive control, with conflict being just one type of such event (Cavanagh et al., 2012; Cavanagh and Frank, 2014). Correlations between theta power and pupil responses suggest that these two signals might reflect activity in neural systems that jointly engage adaptive control processes (Verguts and Notebaert 2009), such as recruiting additional brain systems and adjusting neural gain that are necessary for optimizing decision making.

## 2. Methods

### 2.1. Participants

There were two sessions in this study. During session one, 219 participants (152 females, 64 males, 3 undisclosed; mean age 18.75 ± 1.87 SD) completed an intertemporal choice task online and we fitted the hyperbolic discount function to each participant's choices. We opted to use this discounting function because it has been shown to explain behavioral and neural data very well despite its simplicity (i.e., only one free parameter to be estimated) (Green and Myerson, 2004; van den Bos and McClure, 2013). As with previous work (Kable and Glimcher, 2007), we then invited only participants who were clear hyperbolic discounters to complete session two, the main laboratory experiment when neurophysiological activity was recorded as participants made intertemporal decisions.

68 participants (47 females, 21 males; mean age 18.47 ± 1.56 SD) completed session two, the main experiment. Before data collection, we pre-registered our recruitment procedures (<u>osf.io/7m9c2</u>), which were crucial to our design because we had planned to use each participant's individual hyperbolic function to parametrically vary intertemporal preferences and subjective conflict. All participants provided informed consent in accordance with policies of the university's institutional review board and had normal or correct-to-normal vision.

### 2.2. Experimental design and statistical analysis

Minimum sample size for the main experiment was approximated by running a power analysis in G*Power 3.1 (Faul et al., 2007): For 80% power (*α* = 0.05), 33 participants were required, assuming a repeated-measures ANOVA design with small-to-medium effects (*f* = 0.15), and correlation among repeated measures = .50. Given these estimates, we aimed instead for 60 participants, which gave us roughly 95% power.

The pre-registered design, data, and scripts can be found on the Open Science Framework (<u>osf.io/7m9c2</u>). To summarize, we pre-registered the two-part design that allowed us to model conflict levels for each person separately. Our primary pre-registered prediction was that a response-locked conflict negativity and pupil responses would track conflict in a curvilinear (inverted-U) manner, with largest responses when the subjective values of the two intertemporal rewards were equal (and smaller responses when either of the two rewards had a higher subjective value than the other). We also expected the no-brainer to serve as a control condition with the least conflict. All other analyses and results (i.e., midfrontal theta power, theta-pupil correlations, and increased activity associated with no-brainer choices) are additional and were not pre-registered prior to data collection.

Statistical analyses were performed in R (R Core Team, 2016). The design was entirely within-subjects; unless stated otherwise, all estimates and statistics were obtained by fitting two-level multilevel regression models (all factors and neurophysiological responses for each condition were nested within participants) with random intercepts (unstructured covariance matrix) using the R package lme4 (Bates et al., 2015). When fitting models that tested the relationship between conflict and neurophysiological responses, we included decision time as a predictor of neurophysiological responses. If conflict still significantly predicted these responses after controlling for decision time, the results would suggest that these responses might reflect more than just decision time or choice difficulty. Probability values and degrees of freedom associated with each statistic were determined using the Satterthwaite approximation, using the package lmerTest (Kuznetsova et al., 2016), and an *r* effect size was reported for fixed effects in each model. Bayes factors (BF) were computed by fitting Bayesian multilevel models using the R package brms (Bürkner, 2016).

### 2.3. Stimuli and tasks

During session one, participants completed an online intertemporal choice task where they made 144 decisions. The immediate reward was always $15 today and the delayed reward was $15.50, $24, $42, $71, $107, or $139, offered at a delay of 1, 10, 21, 50, 90, or 180 days (36 unique choice pairs). These values and delays were chosen such that they approximated those used in previous neuroimaging intertemporal choice studies (Kable and Glimcher, 2007; Kable and Glimcher, 2010). Each choice pair (e.g., $15 in 0 days vs. $139 in 10 days) was presented four times. Participants were told that there is no correct answer and they only have to choose they option they prefer. For each participant, we used logistic regression to estimate the indifference point at each delay and non-linear regression to fit the hyperbolic discount function SV = A / (1 + *k*D), where *SV* is subjective value (expressed as the fraction of immediate reward), *A* is delayed reward amount, *D* is delay (in days), and *k* is a participant-specific constant, the only free parameter to be estimated in this model (Green and Myerson, 2004). Each participant has a unique *k* parameter that describes the steepness of the discounting curve, with larger values reflecting steeper slopes. This parameter also captures individual differences in impulsivity, with larger values indicating greater impulsivity or temporal discounting. The hyperbolic discount function for each participant describes how any given delayed reward is translated into a subjective value for that participant. As such, the function provides a principled way of quantifying the subjective value of any delayed reward.

We used each participant's hyperbolic discount function to generate idiosyncratic, participant-specific delayed rewards that allowed us to parametrically vary value difference (i.e., decision conflict) for session two based on the value difference between the immediate and delayed rewards. As with session one, the immediate reward for session two was also always $15 today. The participant-specific delayed rewards (e.g., $24.14 in 10 days) had pre-determined subjective values of 4, 7, 10, 12, 14, 15, 16, 18, 20, 23, and 26 at three different delays (10, 30, and 60 days). We also included three no-brainer, 'catch', delayed rewards that served as control conditions and attention checks, one at each delay: $15 in 10 days, $15 in 30 days, and $15 in 60 days. We expected these no-brainer trials to be the easiest or least conflicting choices. In total, there were 36 unique choice pairs, and each choice pair—including no-brainer choices—was presented with equal probability: three times per block for seven blocks (756 trials in total). On each trial during the EEG and eye tracking session, participants chose between a fixed immediate reward of $15 today and a model-derived participant-specific delayed reward.

During session two, EEG, eye tracking, and electromyography (EMG) data were collected while participants made intertemporal decisions. Participants were told that they are taking part in a decision-making study and that there are no correct or wrong answers, and that they should simply state their preference in a series of choices. They were also told that at the end of the experiment, they might receive one of their choices, which was randomly selected by the computer from the set of all their choices. At the end of the experiment, during debriefing, the experimenter presented each participant with 10 lottery tickets (two are winning tickets). If the participant selected the winning ticket, they received that randomly selected choice, which was paid in the form of an Amazon gift voucher that was emailed to them after the appropriate delay. Given this payment scheme, participants were told at the beginning of the experiment that they should make each choice as though it were the one they are actually going to receive.

Participants completed the task in a dimly-lit room and rested their heads on a chinrest. All stimuli were presented using PsychoPy (Peirce, 2007, 2009) and since the immediate reward was $15 for all trials, the display was simplified to minimize eye movement by presenting only the delayed reward at the center of the screen with a black background (Fig. 1). At a viewing distance of 93 cm, the stimulus (delayed reward) subtended approximately 2.15° horizontally and 0.93° vertically. Participants had up to three seconds to decide whether to choose the immediate reward ($15) or the model-derived delayed reward by pressing either the F or J key (counterbalanced across participants) with their left or right index finger. On each trial, a red central fixation cross appeared for 150 ms. The delayed reward was then displayed in white at the center and remained on screen until the participant responded or until a maximum of 3000 ms had passed. Once the delayed reward had been removed from the screen, a blank black screen appeared for a random interval that varied randomly from 300 to 700 ms. Participants first completed 15 practice trials with feedback showing them what they had chosen ($15 in 0 days or a model-derived delayed reward). They then completed 7 actual blocks of 108 trials each. Each choice pair was presented three times per bock and thus 21 times over 7 blocks. After each block, participants were given the opportunity to rest. They were also told to try to avoid blinking and movement excessively during the experiment. The intertemporal choice task took about 30 minutes to complete.

**Fig. 1.**
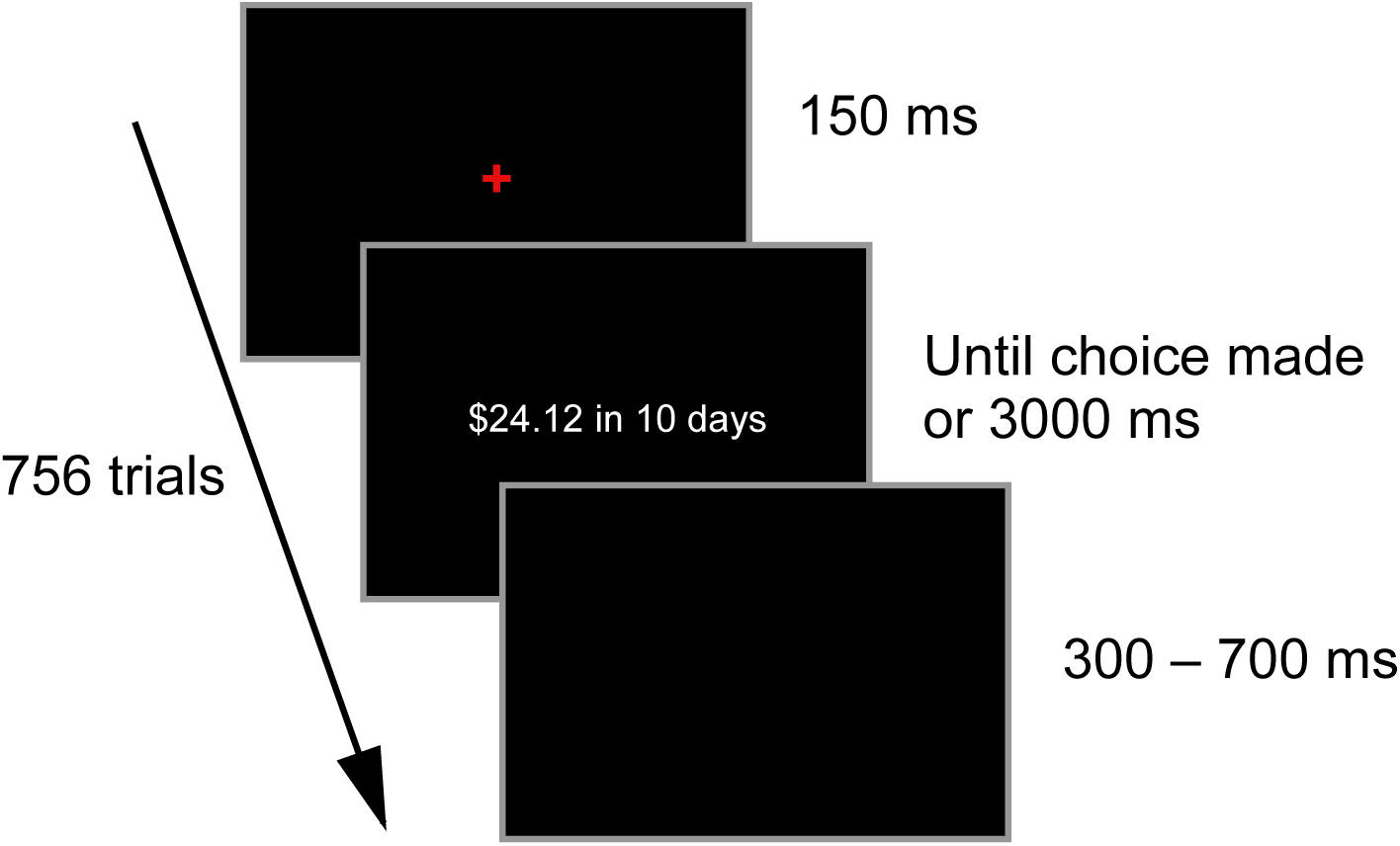
Intertemporal choice task trial sequence. The sequence of events within a trial is shown. On each of 756 trials, participants pressed either the F or J key (counterbalanced across participants) to choose between an immediate ($15 today) and a participant-specific model-derived delayed reward (e.g., $24.14 in 10 days). Participants were told at the beginning of the experiment that the immediate reward was always $15 today and would never be presented visually. Participants had up to 3 s to decide. The inter-trial interval varied randomly from 300 to 700 ms. Seven blocks of 108 choices each were presented. Each of 36 unique choice pairs (including no-brainer choices) was presented three times per block.

### 2.4. Identifying choice outliers

To identify choice outliers, we analyzed behavioral responses for no-brainer catch choices ($15 in 0 days vs. $15 in 10, 30, or 60 days) where we expected participants to always prefer the immediate reward of $15 today. For each participant, we calculated the percentage of no-brainer trials whereby they made the 'wrong' choice (chose $15 in 10, 30, or 60 days rather than $15 today). We then used a robust median absolute deviation outlier detection method to identify participants whose error percentage was three or more median absolute deviations from the median error percentage (Leys et al., 2013). Nine participants made the 'wrong' choice (chose delayed reward) too frequently on no-brainer choices (61.33%; range 26.90−95.00%), and were excluded from all analyses. The remaining 59 participants made 5.50% (range 0−23.81%) 'wrong' choices.

### 2.5. Adjusting subjective values of delayed rewards

Based on the hyperbolic function modeling, we expected participants to experience maximum decision conflict (i.e., theoretical indifference point) when the immediate and delayed rewards have the same subjective values. Theoretically, when both intertemporal options have the same subjective value, participants should choose either intertemporal reward 50% of the time and should also respond slowest. However, previous value-based decision-making studies have shown that it is common to observe discrepancies between theoretical (based on modeling) and empirical (observed behavior) indifference points (Kolling et al., 2012; Kolling et al., 2016; Shenhav et al., 2014). Such discrepancies could lead to incorrect conclusions (Shenhav et al., 2014), especially when the study requires relatively fine differences in subjective values.

We found that our theoretical and empirical indifference points did not perfectly coincide, in that participants had a slight preference for the delayed reward at the theoretical point of indifference (Fig. 2A), suggesting that the subjective value of the delayed rewards were imprecise. Following previous work (Shenhav et al., 2014; Shenhav et al., 2016), we fitted three logistic regressions (one for each intertemporal delay) to each participant's behavioral choices to determine three empirical indifference points at each intertemporal delay, which allowed us to infer from participant's choices the subjective value of each delayed reward. We used the difference between theoretical and empirical difference points to adjust the subjective value of the delayed rewards. The adjusted subjective values would provide more accurate estimates of value difference and decision conflict, and was used as a regressor for all analyses.

**Fig. 2.**
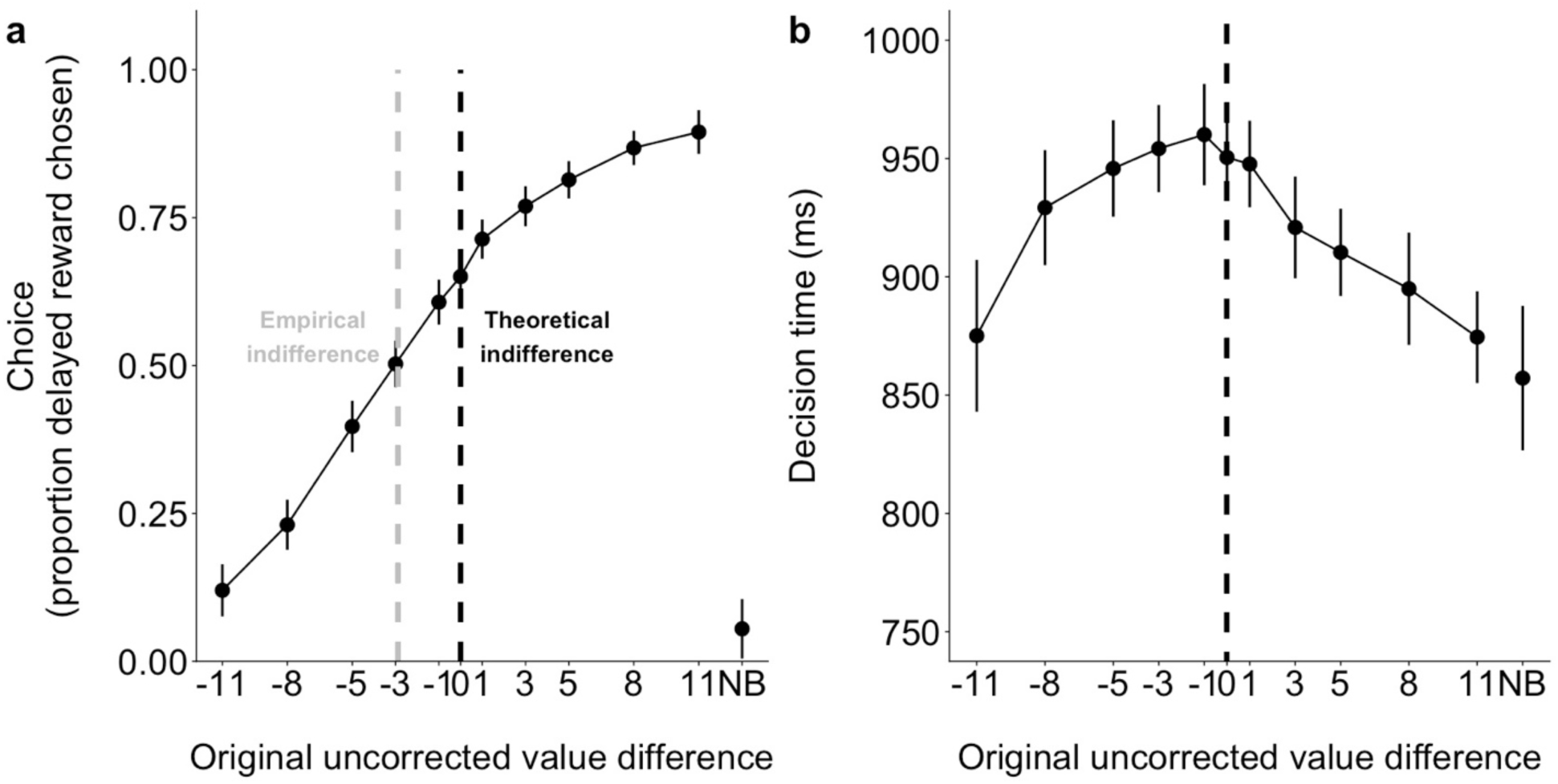
Uncorrected choice proportion and decision time. (A) Sigmoid function for proportion of delayed reward chosen. When difference value < 0, the immediate reward should be chosen more frequently. When difference value > 0, the delayed reward should be preferred. (B) Inverted-U relationship between difference value and decision time. Black dashed line is the theoretical indifference point (model-derived value difference is 0) where participants were expected to be most conflicted. Grey dashed line is the empirical indifference point where participants were choosing as though they were experiencing the most decision conflict. Note the leftward shift of the indifference point. NB refers to no-brainer choice. Error bars indicate 95% confidence intervals.

### 2.6. EEG/EMG recording and preprocessing

Electrophysiological signals were measured via EEG and facial electromyography (EMG) and sampled at 1024 Hz. Impedances were ≤ 5 kΩ during recording. Continuous EEG activity was measured over seven cortical midline sites (Fpz, Fz, FCz, Cz, CPz, Pz, Oz) using Ag/AgCl electrodes embedded in a stretched Lycra cap (Electro-Cap International, Eaton, OH). Because we had not intended to localize neural sources when designing the study and were primarily interested in midfrontal conflict-related activity, we recorded only from midline electrodes and focused primarily on midfrontal sites (i.e., FCz) where conflict activity has typically been observed (Cavanagh et al., 2012; Nakao et al., 2010; Yeung et al., 2004). Vertical electrooculography (VEOG) was recorded around the right eye with two electrodes.

EMG activity over the corrugator supercilii muscles (frowning muscles) were also measured using two electrodes above the inner corner of the left eye, but these data were recorded for other analyses that are unrelated to the present study and will not be reported subsequently^1^. Signals were amplified using ANT TMSi Refa8 device (Advanced Neuro Technology, Enschede, The Netherlands), grounded to the forehead, and referenced online to the average of all electrodes. Offline, EEG signals were re-referenced to the average of electrodes placed on the two earlobes. During pre-processing, continuous data were high-pass filtered at 0.1 Hz (12 dB/oct, zero phase-shift Butterworth filter) and decomposed into independent components using the infomax independent component analysis algorithm implemented in EEGLAB (Delorme and Makeig, 2004). We then used a time-domain approach to identify and remove one independent component^2^ that best correlated (p < .001; via metrics such as correlation and convolution) with eye-blink activity in the VEOG channels (icablinkmetrics toolbox; Pontifex, Miskovic, & Laszlo, 2017).

Time-frequency calculations were computed using custom MATLAB (MathWorks) scripts (Cavanagh et al., 2012; Cohen, 2014a). EEG data were segmented into long epochs of 5000 ms (−2500 to 2500 ms relative to event onset) to avoid potential time-frequency decomposition edge artifacts. Time-frequency measures were computed by multiplying the fast Fourier transformed (FFT) power spectrum of single-trial EEG data with the FFT power spectrum of a set of complex Morlet wavelets, and taking the inverse FFT. The wavelet family is defined as a set of Gaussian-windowed complex sine waves, exp(-i2*πtf*) * exp(-*t*^2^/(2*σ*^2^)), where *t* is time, *f* is frequency (increased from 1 to 30 Hz in 40 linearly spaced steps) and *σ* defines the width (decreased from 0.318 to 0.053) or number of cycles (increased from 4 to 10 in logarithmically spaced steps) of each frequency band. The end result of this process is identical to time-domain signal convolution. Time-frequency power was defined as *Z*(*t*) (power time series: *p*(*t*) = real[*z*(*t*)]^2^ + imag[*z*(*t*)]^2^, and was normalized by conversion to a decibel scale, 10 log_10_ [power(*t*) / power(baseline)], allowing a direct comparison of effects across frequency bands. For stimulus-locked time-frequency power, epochs were baseline normalized for each frequency by the average power from −500 to −200 ms before stimulus onset. Values for statistical analysis were summed over time and frequency (340 to 840 ms, 3.2 to 7.7 Hz), and were based on inspection of the grand average time-frequency power plots (collapsed localizer approach) (Luck and Gaspelin, 2017). For peri-response time-frequency power, epochs were baseline normalized for each frequency by the same pre-stimulus average baseline power described above. Values for statistical analysis were summed over time and frequency (−160 to 40 ms, 3.2−7.7 Hz), and were based on inspection of the grand average time-frequency power plots. To visualize theta power time course for each experimental condition, we also computed theta power over time by computing and plotting mean theta power (3.2−7.7 Hz) at each time point.

To compute event-related potentials (ERPs), pre-processed (0.1 Hz high-pass filtered) EEG signals were low-pass filtered at 20 Hz (12 dB/oct, zero phase-shift Butterworth filter). Epochs were checked for artifacts and automatically rejected using the following criteria: voltage steps of more than 15 µV between sample points, a voltage difference of 150 µV within 150 ms intervals, voltages above 85 µV and below − 85 µV, moving window peak-to-peak voltages exceeding 150 µV (150 ms window with 50 ms step size), and spectra estimates that deviated from baseline by ±50 dB in the 0− 2 Hz frequency window (to detect eye movements) and +25 or −100 dB in the 20−40 Hz frequency window (to detect muscle activity). Mean amplitudes within a selected time window were reported for all ERPs. For stimulus-locked ERPs, epochs were baseline-corrected by the average power from −200 to 0 ms relative to stimulus onset. For response-locked ERPs, epochs were baseline-corrected by the average power from − 200 to −100 ms relative to response onset. Time windows for statistical analysis for stimulus-(500 to 800 ms) and response-locked (0 to 100 ms) ERPs were determined based on inspection of the grand-average ERP waveforms (Luck and Gaspelin 2017). Because the stimulus-locked N2 component has typically been associated with response conflict during inhibitory control tasks (Yeung et al., 2004), we also visually inspected the grand-average ERP waveform to localize the second negativity after stimulus onset (i.e., 340 to 440 ms).

### 2.7. Eye tracking and pupil dilation preprocessing

Pupillometric data were recorded using the EyeLink 1000 Desktop Mount eye tracker (SR Research, Mississauga, Ontario, CA). The EyeLink system was configured using a 35-mm lens, 5-point gaze location calibration, monocular right-eye sampling at a rate of 1000 Hz, and centroid fitting for pupil area recordings. Pupil measures reflected pupil area. All data processing were performed using custom R scripts. Blink artifacts detected using the EyeLink blink detection algorithm were removed using linear interpolation from 100 ms prior to and 200 ms post blink onset (Cavanagh et al., 2014). Because blinks usually do not last longer than 500 ms, any time window where pupil data were missing for ≥ 500 ms was not interpolated (blinks do not typically last longer than this duration) and instead treated as missing data.

Continuous data were epoched (−500 to 4000 ms) surrounding the onset of stimulus. Trials with decision times ≤ 250 ms or above the decision time deadline of 3000 ms were excluded. Pupil responses were calculated as the percent change from the trial-specific pre-fixation baseline mean (−500 to −300 ms: only a black blank screen was shown). Stimulus-induced pupil dilation responses begin with a light-induced constriction and recovery that last for about 1000 ms, and pupil dilations are very slow and thus are lagged in time to eliciting events, often peaking about 1000 ms (Cavanagh et al., 2014; van Steenbergen and Band, 2013).

To determine the time window where changes in mean pupil dilation response were curvilinearly associated with value difference (statistically significant quadratic coefficient), we fitted a quadratic model at each millisecond from 1001 to 3000 ms after stimulus presentation and controlled for error rates with false discovery rate (FDR) correction (Benjamini and Hochberg, 1995). We used this data-driven mass univariate approach because pupil dilation responses are much more protracted than neural responses and it can be difficult to visually determine the temporal dynamics of an effect. The first 1000 ms was excluded from this analysis because stimulus-induced pupil dilation responses begin with a light-induced constriction and recovery that last for about 1000 ms, and pupil responses are slow and thus are lagged in time to eliciting events, often peaking after about 1000 ms (Cavanagh et al., 2014; van Steenbergen and Band, 2013). The time window from 1110 to 3000 ms survived FDR correction (p *<* .05) and was used to calculate the mean pupil dilation response associated with each experimental condition.

### 2.8. Theta-pupil correlations over time

To integrate midfrontal theta and pupil data, we first used the region-of-interest method by correlating across all participants mean theta power (3.2−7.7 Hz from 340 to 840 ms) with mean pupil data (1001 to 3000 ms) associated with each experimental condition computed for each participant separately. To further probe theta-pupil relationships over time, we explored correlations across the entire time series (rather than just region-of-interest) for EEG and pupil data (Chmielewski et al., 2017). We downsampled both the EEG and pupil data to 50 Hz (each time point is 20 ms), and then computed the correlation coefficient across all participants, using each time point from the entire EEG theta series (−500 to 1500 ms) with each time point from the entire pupil time series (−500 to 3000 ms). 17776 correlations were performed, and we used FDR correction to control for error rates, and visualized only correlations that survived FDR correction (p < .05).

## 3. Results

### 3.1. Choice and reaction time reflect subjective value comparison and conflict

Choice and decision time patterns suggest that participants had compared the subjective values of the immediate and delayed rewards and experienced subjective decision conflict (Fig. 3). When the immediate and delayed rewards were equally desirable (value difference = 0), participants chose the delayed reward 51% of the time (Fig. 3A). A logistic regression indicated that choice was predicted by value difference, in that participants were more likely to choose the delayed over the immediate reward (coded as 1 and 0 respectively) as value difference increased from −11 to +11 (b = 0.28, z = 91.08, p < .001). As predicted, the relationship between value difference and decision time was curvilinear (quadratic b = −0.13, SE = 0.009, t(1178) = −14.71, p < .001, r = .39), suggesting that participants could discern relatively fine differences in subjective value (Fig. 3B): Decision time was slowest (mean = 1055 ms; SD = 142 ms) when the immediate and delayed rewards had the same subjective value (value difference = 0); they were much faster when the immediate reward was clearly better than the delayed reward (value difference = −11; mean = 936 ms; SD = 151 ms) or vice versa (value difference = +11; mean = 847 ms; SD = 102 ms). Decision times for no-brainer choices were also relatively fast (mean = 842 ms; SD = 134 ms; Fig. 3B), suggesting little subjective conflict.

**Fig. 3.**
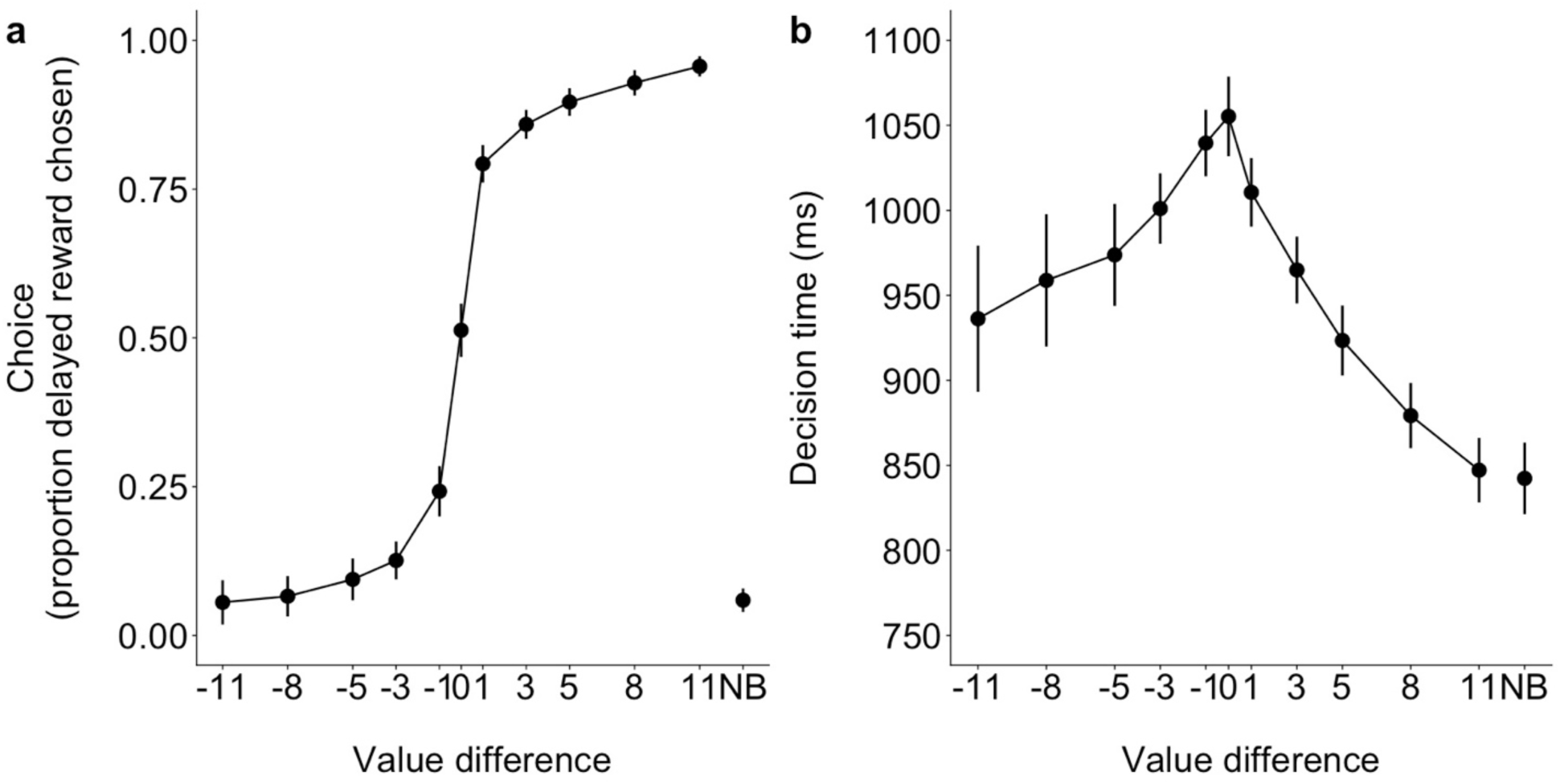
Choice proportion and decision time based on adjusted value difference. Logistic regression was used to adjust subjective values for each participant based on observed choice. (A) Sigmoid function for proportion of delayed reward chosen. When value difference was 0, the immediate and delayed rewards had the same subjective value of 15. When difference values were negative, the delayed rewards had smaller subjective values relative to the immediate reward and were chosen less frequently. When difference values were positive, delayed rewards had larger subjective values and were chosen more frequently. (B) Inverted-U relationship between value difference and decision time. NB refers to the no-brainer choice. Error bars indicate 95% confidence intervals.

### 3.2. EEG dynamics: Midfrontal theta power and event-related potentials (ERPs)

We observed a robust increase in stimulus-locked theta power (3.2−7.7 Hz) over midfrontal scalp electrodes after choice presentation. Because previous studies have observed strongest conflict effects on electrode FCz (e.g., Cavanagh et al., 2012), we also focus our analyses on this electrode. Theta power (340 to 840 ms after choice onset) was enhanced when the immediate and delayed rewards had similar subjective values (Fig. 4A). As predicted, value difference was curvilinearly related to theta power (quadratic b = −0.26, SE = 0.05, t(1181) = −5.60, p < .001, r = .16), indicating that theta power decreased as it became increasingly clearer that one reward had a higher subjective value than the other (Fig. 4C). Additional analyses suggested that this effect was maximal at electrode FCz: Despite the lack of interaction between electrode site (frontal: FPz, Fz; central: FCz, Cz, CPz; posterior: Pz, Oz) and quadratic coefficient (F(2, 8573) = 0.36, p = .699), we found that the effect was strongest at the FCz site (b = −0.26, p < .001, r = .16), weaker at frontal sites (FPz: b = −0.16, p = .025, r = .07; Fz: b = 0.25, p = .339, r = .03), central sites (Cz: b = −0.20, p < .001, r = .13; CPz: b = −0.12, p = .007, r = .08), and posterior sites (Pz: b = −0.06, p = .155, r = .04; Oz: b = 0.01, p = .792, r = .01). We also found that the curvilinear relationship between value difference and midfrontal theta power was driven primarily by non-phase-locked theta power (quadratic b = −0.39, SE = 0.07, t(1185) = −5.56, p < .001, r = .16), rather than phase-locked theta power (quadratic b = 0.03, SE = 0.01, t(1191) = 2.33, p = .020, r = .07), suggesting that the effects we have observed might be more apparent in the time-frequency rather than time-domain (i.e., ERPs). For details and figures related to these analyses, refer to the Supplementary Material.

Although the inverted-U relationship between value difference and theta power appears to suggest that midfrontal theta parametrically subjective conflict (Fig. 4), three additional findings indicate that theta power reflects more than just variation in decision conflict. First, a conflict account would predict that no-brainer choices should be the least conflicting of all choices and should therefore evoke minimal theta power. However, unexpectedly, theta power for no-brainer choices was significantly and robustly higher than the mean theta power of all other choices combined (b = 0.35, SE = 0.06, t(1348) = 6.11, p < .001, r = .16; Fig. 4). Moreover, theta power for no-brainer choices was not significantly different from that associated with the most conflicting choices whereby value difference is 0 (b = −0.14, SE = 0.09, t(292) = −1.55, p = .122, r = .09). To supplement this null finding, we computed a Bayes Factor (BF) to test two hypotheses (H0: theta_no-brainer_ = theta_most-conflict_; H1: theta_no-brainer_ < theta_most-conflict_), and found moderate evidence favoring the hypothesis that theta power for no-brainer and the most conflicting choices were in fact equal (BF01 = 3.86), despite no-brainer choices being an easy decision. Because this finding was unexpected, we ran additional analyses and found that no-brainer decisions were also associated with increased delta power in posterior regions (see Supplementary Material). Second, theta power for all choices was stronger when delayed rewards were presented sooner (10 days) rather than later (30 or 60 days) (b = −0.002, SE = 0.001, t(1195) = −2.41, p = .016, r = .07), suggesting that theta power also tracks the immediacy or saliency of the delayed reward. Third, we included decision time in the model but it did not predict theta power (b = −0.04, SE = 0.13, t(480) = −0.33, p = .741, r = .02), and all results were similar after controlling for decision time (quadratic b = −0.27, SE = 0.05, t(1227) = −5.37, p < .001, r = .15), suggesting that theta power reflected more than just decision time or choice difficulty. Taken together, these three findings suggest that midfrontal theta might indicate the need to increase attention and engage adaptive control, rather than track conflict *per se*.

**Fig. 4.**
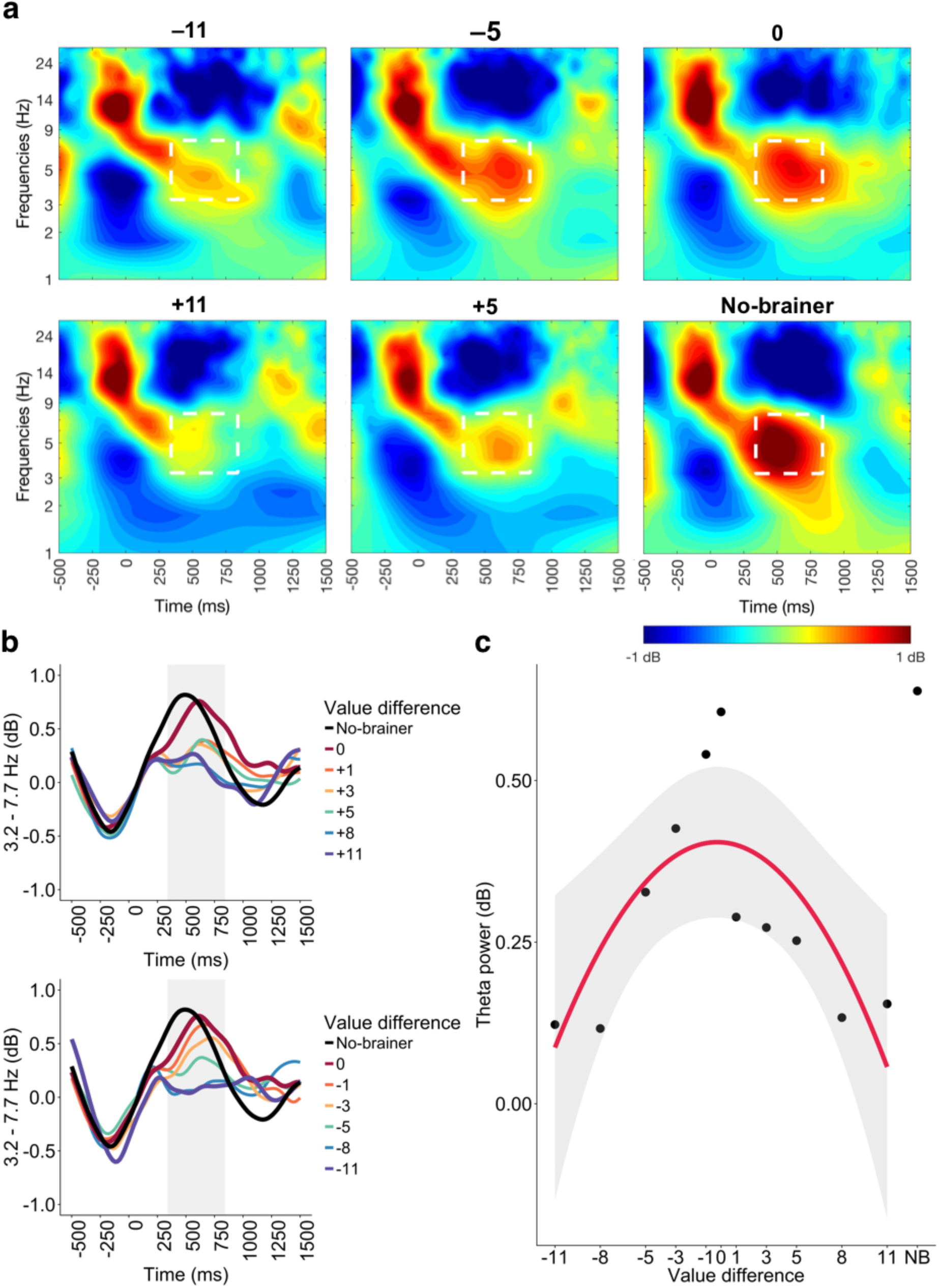
Theta power enhancement after stimulus presentation over midfrontal scalp electrode (FCz), collapsed over different intertemporal delays (10, 30, and 60 days). Increased power (3.2−7.7 Hz) was observed between 340 to 840 ms. (A) Theta power for selected value differences of −11, −5, 0, +5, +11, and no-brainer choice. White-dashed box shows region of interest used to compute mean theta power. (B) Theta power (3.2−7.7 Hz) time course. Top and bottom panels show positive and negative value differences respectively. (C) Curvilinear relationship between value difference and theta power. NB refers to no-brainer choice. Error bars indicate 95% confidence intervals.

Since previous studies have often reported conflict-related ERPs, we also expected to subjective conflict to modulate stimulus-locked ERPs at midfrontal regions (FCz electrode), specifically the N2. However, unlike previous studies on objective response conflict using inhibitory control tasks (Yeung et al., 2004), subjective conflict in our study did not modulate the N2 component (340 to 440 ms; quadratic b = 0.12, SE = 0.20, t(1179) = 0.62, p = .538, r = .02), but instead modulated a positive-polarity ERP over the FCz electrode from 500 to 800 ms after choice onset (Fig. 5A). Value difference was curvilinearly related to the amplitude of this ERP (Fig. 5B; quadratic b = −0.51, SE = 0.20, t(1220) = −2.50, p = .013, r = .07), suggesting that this ERP also parametrically tracked subjective conflict during intertemporal choice (but the effect was only modest in size). The amplitude of this ERP was associated with decision time (b = −1.23, SE = 0.56, t(856) = −2.20, p = .028, r = .07), but not whether delayed rewards were presented sooner or earlier (b = −0.004, SE = 0.004, t(1192) = −1.22, p = .222, r = .04). As with theta power, this ERP also appeared to track more than just conflict. The amplitude of this ERP for no-brainer choices was larger than that associated with all the other choices combined (b = 1.17, SE = 0.23, t(1349) = 5.10, p < .001, r = .14; Fig. 5B). The amplitude for no-brainer choices did not significantly differ from that associated with the most conflicting choices (b = 0.31, SE = 0.35, t(288) = 0.87, p = .386, r = .05), and a Bayesian analysis found strong evidence favoring the hypothesis that ERP amplitudes for these two types of choices were in fact equal (BF01 = 54.88).

**Fig. 5.**
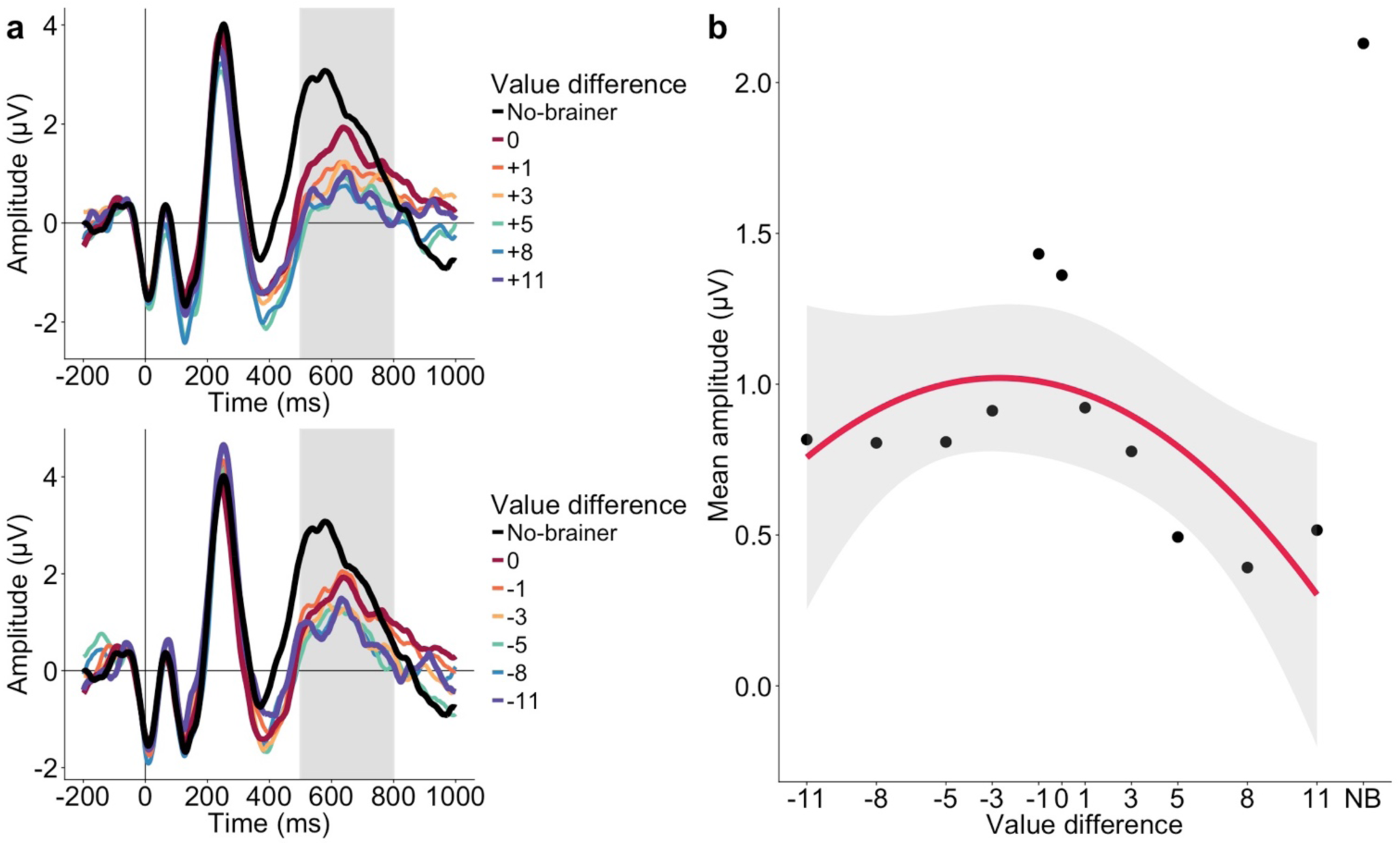
Event-related potential after stimulus presentation at FCz electrode. (A) A positivity resembling the P300 was observed around 500−800 ms. (B) Mean amplitude of positivity for each value difference and no-brainer choice. NB refers to no-brainer choice. Error bars indicate 95% confidence intervals.

Because previous work on conflict using inhibitory control tasks also found that response-locked midfrontal signals tracked binary high versus low conflict (e.g., Cavanagh et al., 2012), we therefore predicted in our pre-registration that these response-locked conflict signals would track different levels of subjective conflict. Indeed, consistent with previous work, peri-response (−360 to 140 ms relative to response onset) theta power (3.2−7.7 Hz) tracked subjective conflict during intertemporal decisions (Fig. 6A). Value difference was curvilinearly related with theta power (b = −0.22, SE = 0.05, t(1183) = −4.54, p < .001, r = .13; Fig. 6C). As with stimulus-locked theta power, peri-response theta power was stronger when delayed rewards were presented sooner (10 days) rather than later (30 or 60 days) b = −0.002, SE = 0.001, t(1204) = −2.22, p = .027, r = .06), suggesting that theta power might track the immediacy or saliency of the delayed reward. After including decision time (b = 0.30, SE = 0.13, t(374) = 2.39, p = .017, r = .12), the curvilinear relationship between theta power and value difference remained significant (b = −0.18, SE = 0.05, t(1228) = −3.51, p < .001, r = .10), suggesting that theta power reflected more than just decision time or choice difficulty. Peri-response theta power for no-brainer choices was significantly and robustly higher than the mean theta power of all other choices combined (b = 0.36, SE = 0.06, t(1351) = 6.35, p < .001, r = .17; Fig. 6C), suggesting that midfrontal theta tracks more than just conflict. Moreover, theta power for no-brainer choices was not significantly different from that associated with the most conflicted choices whereby value difference is 0 (b = −0.03, SE = 0.09, t(293) = −0.36, p = .718, r = .02); a Bayesian analysis also found strong evidence favoring the hypothesis that theta power for no-brainer and the most conflicting choices were equal (BF01 = 11.58). These findings suggest that midfrontal theta might track more than just conflict, and may instead reflect increased attentional and adaptive control.

**Fig. 6.**
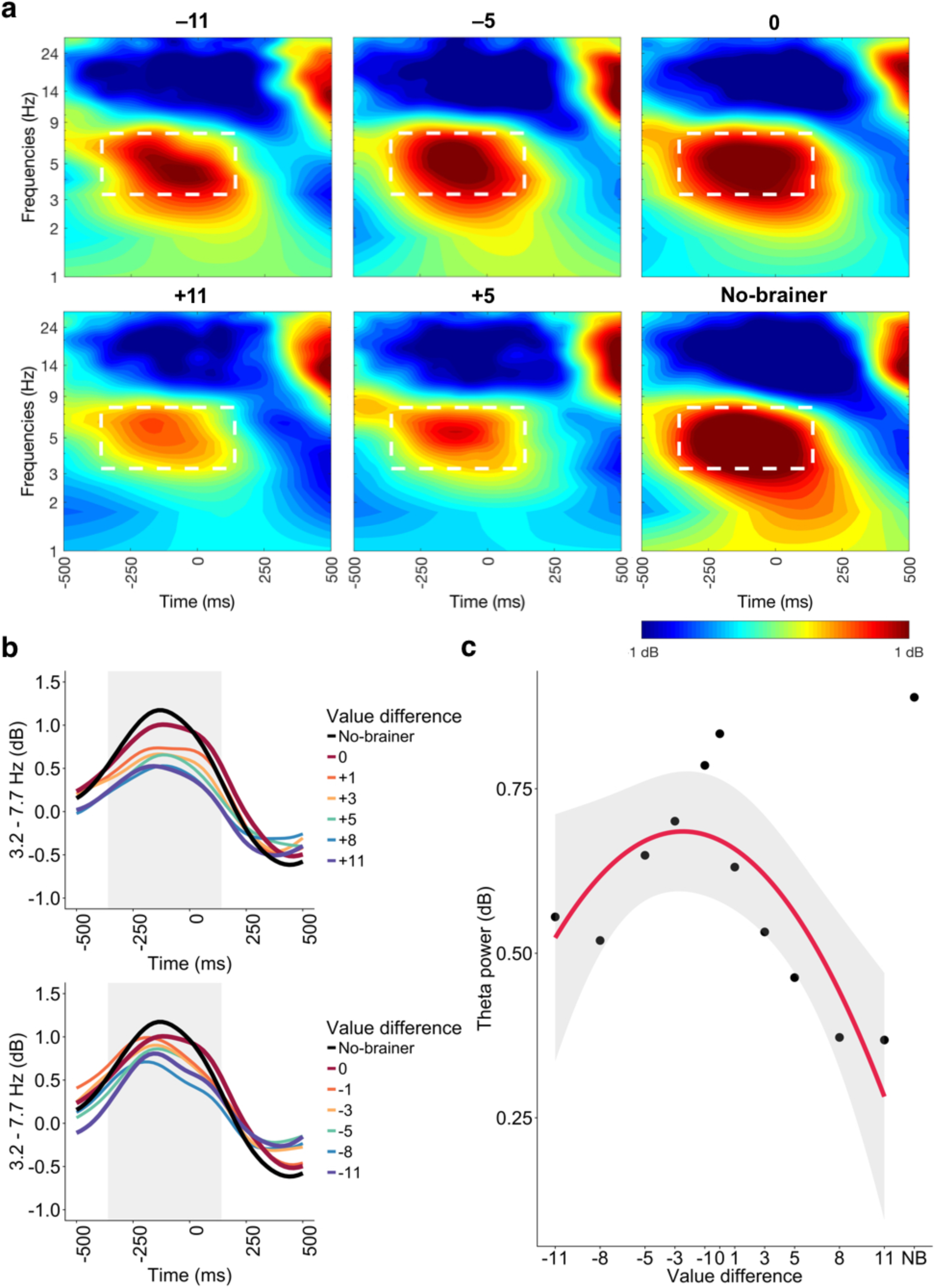
Theta power enhancement after around response over midfrontal scalp electrode (FCz), collapsed over different intertemporal delays (10, 30, and 60 days). Increased power (3.2−7.7 Hz) was observed between −360 to 140 ms around response onset. (A) Theta power for selected value differences of −11, −5, 0, +5, +11, and no-brainer choice. White-dashed box shows region of interest used to compute mean theta power. (B) Theta power (3.2−7.7 Hz) time course. Top and bottom panels show positive and negative value differences respectively. (C) Curvilinear relationship between value difference and theta power. NB refers to no-brainer choice. Error bars indicate 95% confidence intervals.

Finally, in a similar peri-response window (0 to 50 ms), we also observed a peri-response ERP that was parametrically modulated by subjective conflict during intertemporal choice (Fig. 7A). This ERP resembled neural responses typically observed following correct responses during inhibitory control tasks (Vidal et al., 2000), as well as the conflict negativity that has been shown to track binary high versus low subjective conflict (Di Domenico et al., 2016; Nakao et al., 2010). Value difference was curvilinearly associated with the amplitude of this ERP (quadratic b = 0.76, SE = 0.21, t(1182) = 3.58, p < .001, r = .10), but whether delayed rewards were presented sooner or earlier was not associated with amplitude, (b = −0.001, SE = 0.004, t(1197) = −0.30, p = .766, r = .10). After including decision time (b = −0.94, SE = 0.58, t(538) = −1.61, p = .109, r = .07), the quadratic relationship remained significant (quadratic b = 0.63, SE = 0.23, t(1227) = 2.82, p = .005, r = .08). Unlike the other EEG responses reported earlier, the amplitude of this ERP for no-brainer choices was not larger than that associated with all the other choices combined (b = −0.42, SE = 0.26, t(1356) = −1.63, p = .102, r = .04; Fig. 7). These findings therefore suggest that subjective conflict during intertemporal choice parametrically modulates this peri-response negativity.

**Fig. 7.**
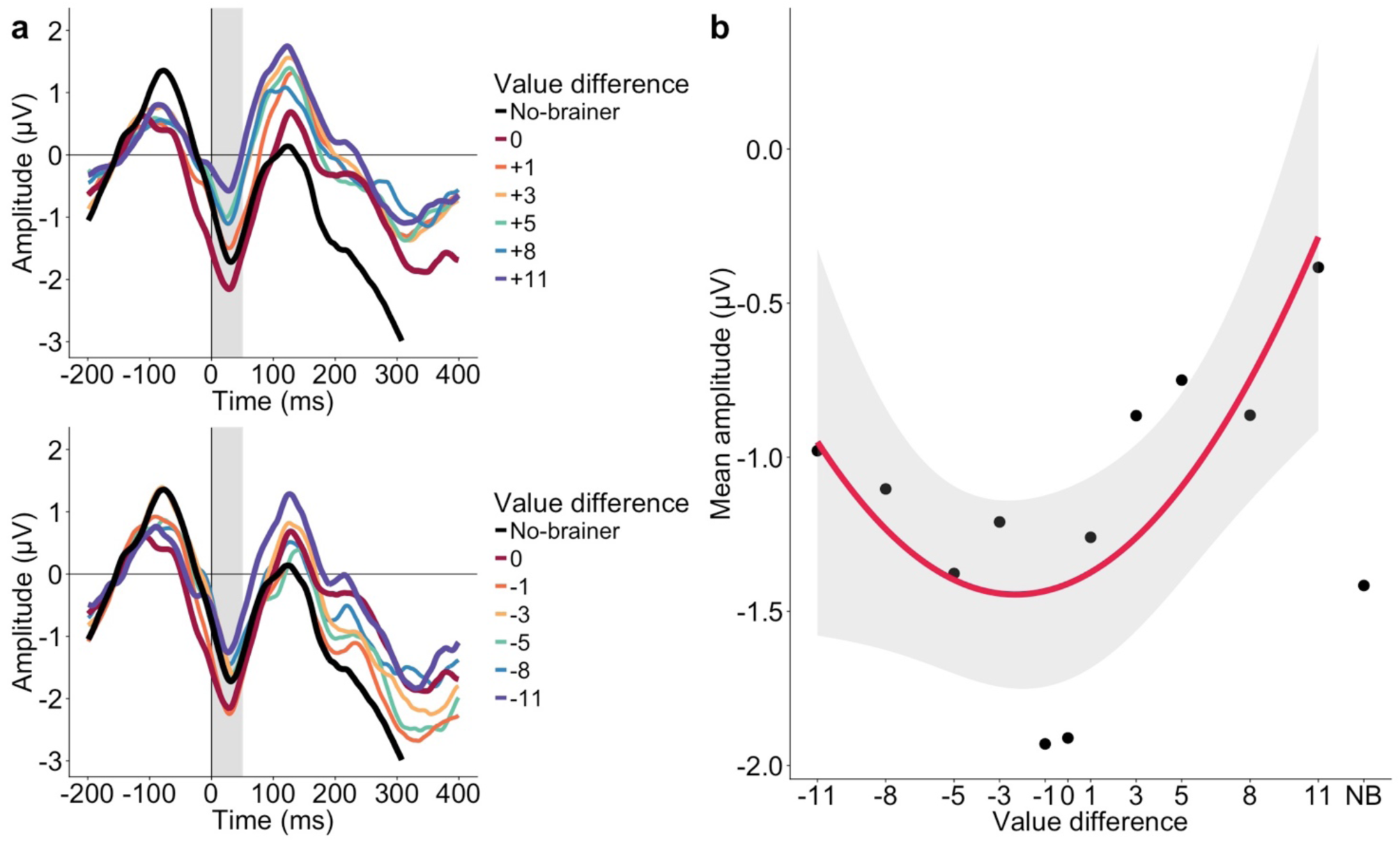
Event-related potential around response onset at FCz electrode. (A) The conflict negativity was observed around 0−50 ms. (B) Mean amplitude of conflict negativity for each value difference and no-brainer choice. NB refers to no-brainer choice. Error bars indicate 95% confidence intervals.

In summary, we found that having competing subjective preferences for different intertemporal rewards induces subjective conflict, which parametrically modulated stimulus-and response-locked theta power (3.2−7.7 Hz) and ERPs measured over midfrontal scalp electrodes (FCz). Unexpectedly, the surprisingly easy no-brainer choices also evoked enhanced midfrontal theta, and theta power was also enhanced when delayed rewards were presented sooner rather than later. These findings suggest that midfrontal theta power do not track conflict *per se*, but might track motivationally relevant events that require attending to, such as high conflict or surprisingly easy no-brainer choices. Subjective conflict in our value-guided choice paradigm did not modulate the N2 component that is usually associated with conflict effects (Yeung et al., 2004); instead, subjective conflict modulated a stimulus-locked positivity (Fig. 5) that resembled the early P3 component (O’Connell et al., 2012; Twomey et al., 2015), which has been associated with decisions processes that reflect phasic LC-NE system activity (Nieuwenhuis et al., 2005; Nieuwenhuis et al., 2011). In addition, the response-locked conflict negativity was the only signal that tracked just subjective conflict but not the immediacy of the delayed reward or the surprising no-brainer choices (Fig. 7). The N2, P3, and conflict negativity findings suggest that conflict and adaptive control processes might be reflected in different ERPs during different types of decisions (e.g., inhibitory control, value-based choice).

Taken together, our subjective conflict results suggest much consilience across different types of decisions (e.g., inhibitory control, reinforcement learning, value-guided decisions) and EEG signals (e.g., theta power, ERPs), but the finding that no-brainer choices also increase midfrontal theta power suggests that more work needs to be done to understand the functional relevance of midfrontal ERPs and theta oscillations.

### 3.3. Pupil dilation responses

We found that subjective conflict parametrically modulated post-stimulus pupil dilation responses (Fig. 8). Value difference was curvilinearly related to pupil dilation responses (quadratic b = −0.64, SE = 0.14, t(1137) = −4.47, p < .001, r = .13), indicating that pupil responses decreased as one reward became more desirable than the other. Pupil responses were not influenced by whether delayed rewards were presented sooner or later (b = −0.005, SE = 0.003, t(1158) = −1.58, p = .114, r = .05). When decision time was included (b = 0.76, SE = 0.38, t(313) = 2.02, p = .044, r = .11), all the above effects remained significant (quadratic b = −0.54, SE = 0.15, t(1179) = −3.58, p < .001, r = .10).

As with EEG signals, pupil responses seem to track more than just conflict. Pupil responses for no-brainer choices were larger than that associated with all the other choices combined (quadratic b = 0.67, SE = 0.17, t(1296) = 3.93, p < .001, r = .11). Pupil responses for no-brainer choices and the most conflicting choices did not significantly differ (b = −0.05, SE = 0.25, t(280) = −0.20, p = .842, r = .01), and there is strong evidence favoring the hypothesis that pupil responses for these two types of choices were equal (BF01 = 36.76).

**Fig. 8.**
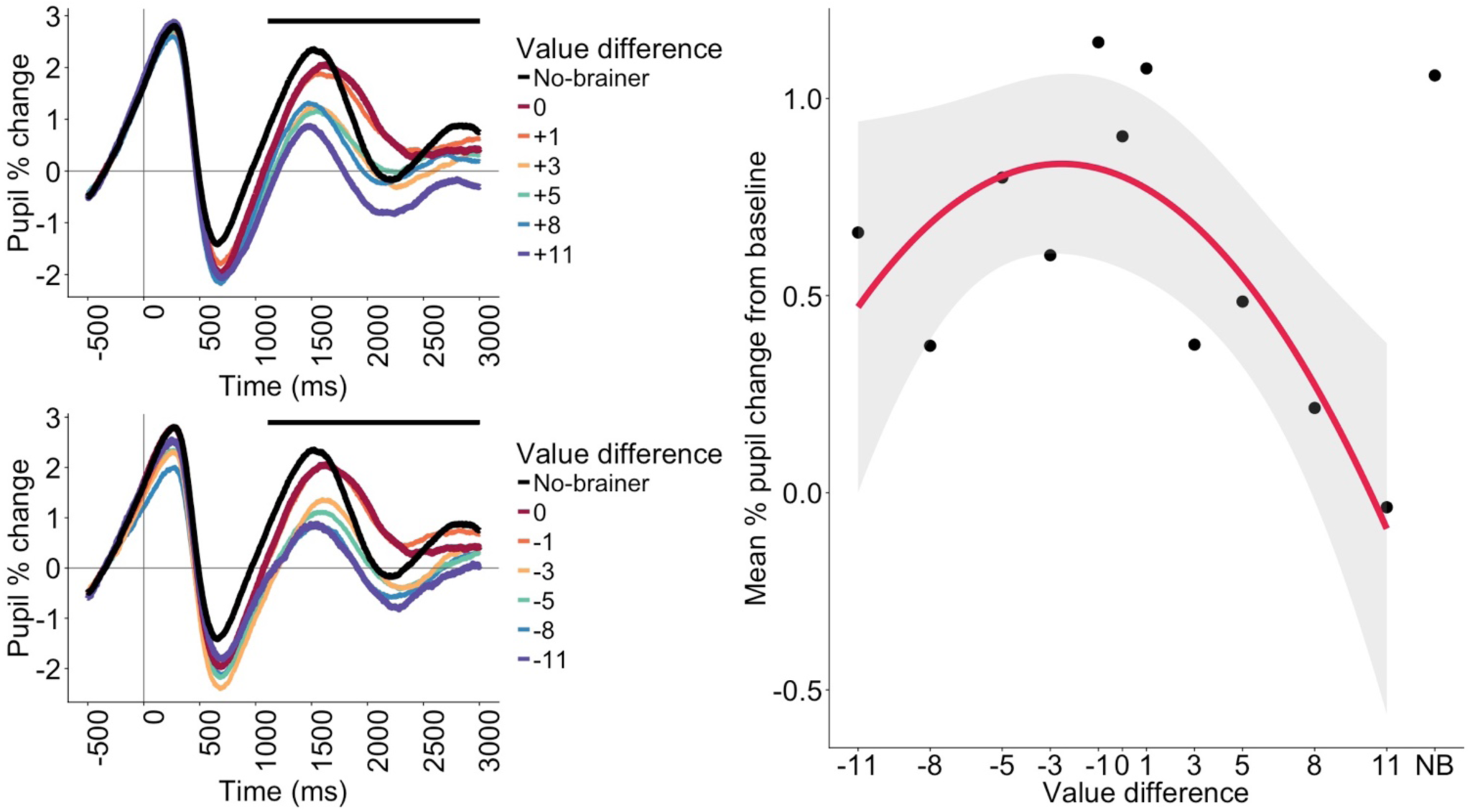
Pupil dilation response after stimulus presentation. (A) Pupil dilation responses tracked value difference. Black horizontal line shows time points (1110 to 3000 ms) that survived FDR correction (p < .05). (B) Mean pupil dilation response (1110 to 3000 ms) for each value difference and no-brainer choice. NB refers to no-brainer choice. Error bars indicate 95% confidence intervals.

### 3.4. Pupil responses associated with EEG dynamics

We performed within-person multi-level model analysis to integrate midfrontal theta and pupil data. We correlated theta power (mean power: 3.2−7.7 Hz, 340 to 840 ms) with pupil dilation (mean pupil size: 1001 to 3000 ms) associated with each experimental condition computed for each participant separately. Midfrontal theta power correlated with pupil dilation responses, (b = 0.03, SE = 0.009, t(1330) = 3.76, p < .001, r = .10, BIC = 2870). Although this effect was small, it was robust and remained significant after controlling for decision time and subjective conflict, (b = 0.03, SE = 0.009, t(1333) = 3.22, p = .001, r = .09, BIC = 2880), suggesting that the theta-pupil relationship was not driven by subjective conflict or decision time. Pupil responses also correlated with the positive-polarity stimulus-locked ERP (b = 0.11, SE = 0.04, t(1313) = 3.07, p = .002, r = .08, BIC = 6591), but the effect became non-significant after including subjective conflict (value difference) in the model (b = 0.03, SE = 0.04, t(1158) = 0.83, p = .407, r = .02, BIC = 5833), suggesting that it might be subjective conflict that is driving the relationship between pupil responses and this ERP.

To further explore the theta-pupil relationship, we correlated 20 ms time bins from the mean theta power (3.2−7.7 Hz) and pupil responses time series (Fig. 9). Soon after choice onset, theta power at 250 to 750 ms significantly correlated with pupil responses from 750 to 1500 ms. These correlations suggest that during value-guided choice, the neural systems thought to underlie midfrontal theta power and pupil dilation responses might interact soon after choice onset.

Although our methods were correlational, we attempted to address directionality effects by fitting basic Granger causality models^3^ on the grand average midfrontal theta power and pupil dilation time courses (−500 to 1500 ms), with lags ranging from 1 to 30. These analyses suggest that midfrontal theta power might be causing pupil dilation responses (ps < .05 for all 30 models that used theta power to predict pupil dilation), rather than pupil dilation responses driving midfrontal theta power (ps < .05 for only 16 of the 30 models that used pupil dilation to predict theta power).

**Fig. 9.**
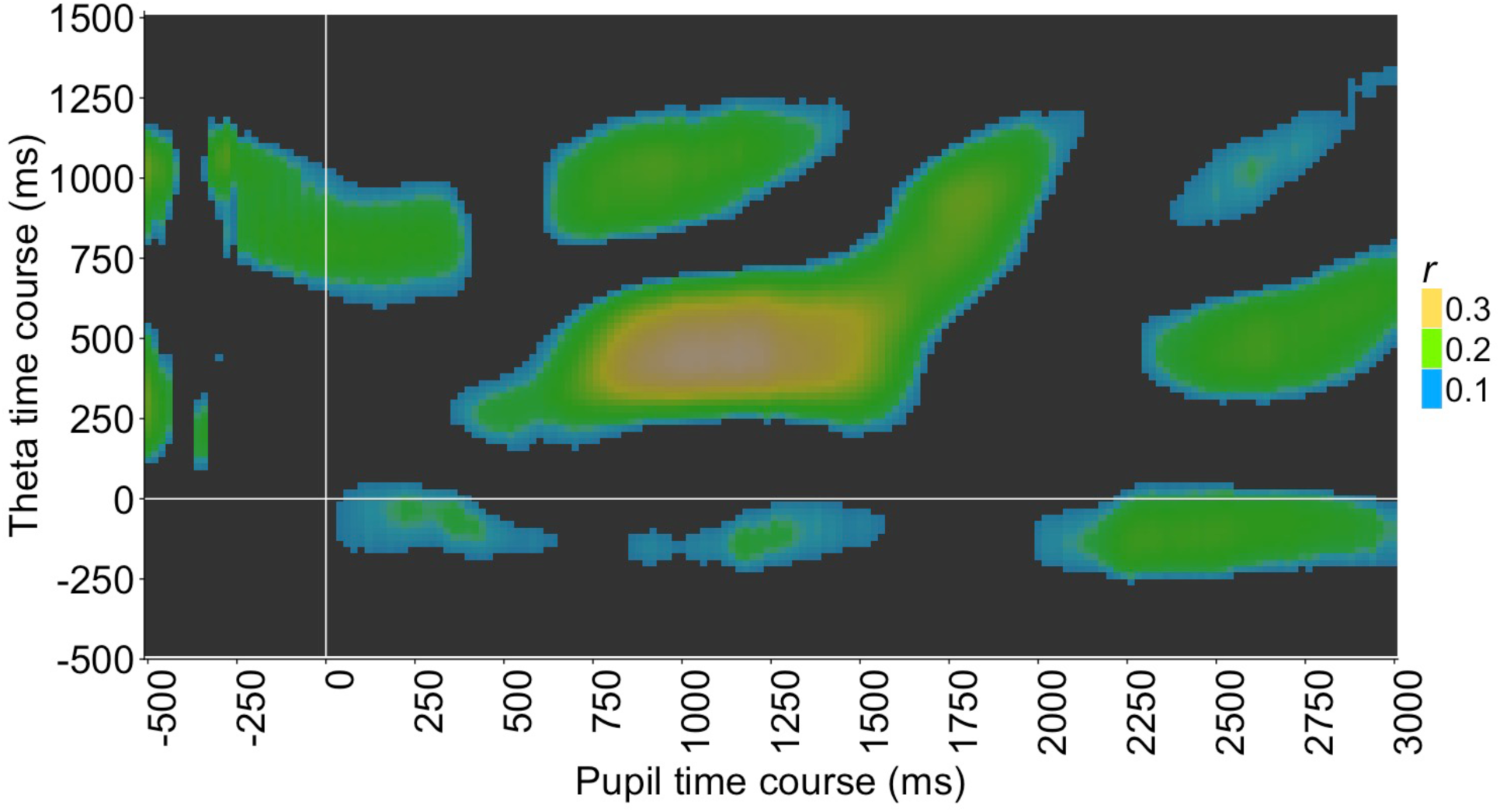
Theta-pupil correlations across entire time series. The x-axis reflects time points (in 20 ms bins) across pupil dilation time series (−500 to 3000 ms), whereas the y-axis reflects time points (in 20 ms bins) across theta power (3.2−7.7 Hz) time series (−500 to 1500 ms). Intertemporal choices were presented at 0 ms. Each time point in each series was correlated with time points in the other series, and the strength of the relationship at each time point is reflected in the heatmap. Colored (non-black) regions survived FDR correction (p < .05), with brighter colors indicating stronger correlations between midfrontal theta power and pupil dilation responses. Strongest correlations were observed during these periods: midfrontal theta (250 to 750 ms) and pupil responses (750 to 1500 ms).

### 3.5. Midfrontal theta power and pupil dilation responses predict decision time

Given that midfrontal theta power, ERPs, and pupil dilation were correlated, we ran additional analyses to explore which stimulus-locked neurophysiological response best predicts behavior (i.e., decision time). First, we used midfrontal theta power and the positive-polarity ERP to predict decision time in the same model. Decision time was significantly associated with midfrontal theta power (b = 0.03, SE = 0.006, t(1189) = 4.31, p < .001, r = .12) but not the ERP (b = −0.001, SE = 0.001, t(1199) = −0.66, p = .511, r = .02). Next, we included pupil dilation responses in the model, which allows us to estimate the effect of each of the three neurophysiological responses on behavior while controlling for the other two responses. The relationship between decision time and the positive-polarity ERP remained non-significant (p = .390, r = .03), but both midfrontal theta power (b = 0.02, SE = 0.006, t(1140) = 3.22, p = .001, r = .10) and pupil dilation responses (b = 0.01, SE = 0.002, t(1137) = 5.81, p < .001, r = .17) were associated with decision time. The effect sizes suggest that when these three neurophysiological measures are included in the same model, pupil dilation responses appear to be the best predictor of decision time.

We then ran mediation analysis to explore whether the stimulus-locked neurophysiological responses (midfrontal theta power, positive-polarity ERP, pupil dilation responses) mediate the relationship between value difference (subjective conflict) and decision time. These three neurophysiological responses were entered as mediators into the same mediation model. This analysis was tested using a bootstrap procedure implemented with the R package “mediation” (Preacher and Hayes, 2008; Tingley et al., 2014). 5000 bootstrap resamples were generated to estimate the sizes and standard errors of the direct, indirect, and total effects. 95% confidence intervals were determined from the bootstrap resamples and any interval that did not include 0 was considered to be significantly different from 0. This mediation analysis (Fig. 10) suggests that the relationship between value difference and decision time is mediated by midfrontal theta power (a_1_b_1_ = 0.02, SE = 0.005, 95% CI [0.01, 0.03], p < .001), and pupil dilation responses (a_2_b_2_ = 0.01, SE = 0.004, 95% CI [0.01, 0.02], p < .001), but not the positive-polarity ERP (a_3_b_3_ = 0.005, SE = 0.003, 95% CI [−0.002, 0.01], p = .15). Overall, the total indirect (mediation) effect for the set of three mediators was significant (f = 0.04, SE = 0.01, 95% CI [0.03, 0.05]). The direct effect of value difference on decision time remained significant after including these three mediators (c’ = −0.16, SE = 0.02, 95% CI [−0.20, −0.12]), suggesting that these three neurophysiological responses only partially mediated the relationship between value difference (subjective conflict) and decision time.

**Fig. 10.**
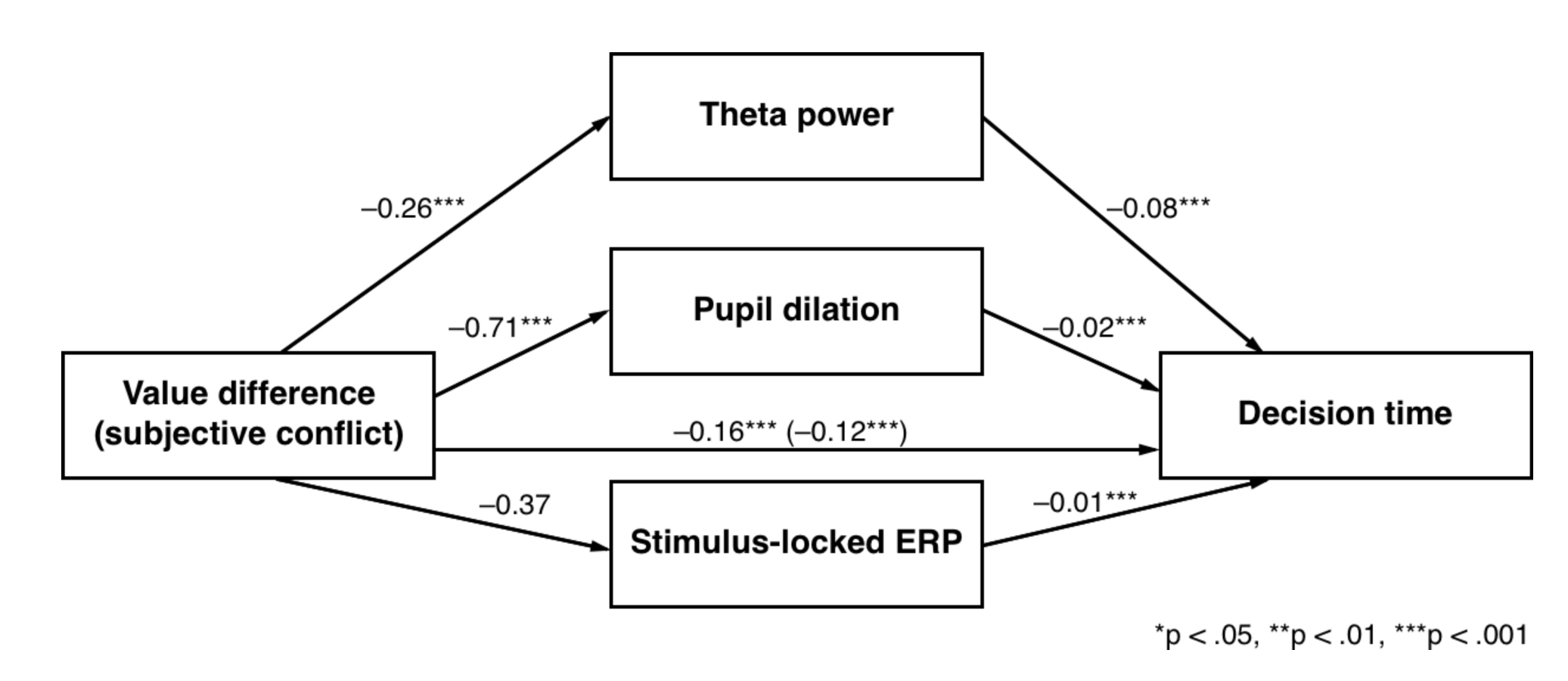
Multiple mediation model of midfrontal theta, stimulus-locked ERP, and pupil dilation responses as mediators of the relationship between value difference (subjective conflict) and decision time. Unstandardized regression coefficients are provided along the paths.

### 3.6. No-brainer choices might be associated with different decision strategies

Given that no-brainer choices elicited neurophysiological responses that were larger than expected, we ran exploratory analyses on decision times, which might provide insights into the types of decision strategies participants were relying on during no-brainer trials. On no-brainer trials, participants had to choose between either $15 today or $15 in 10, 30, or 60 days. We used each participant’s hyperbolic function to determine the subjective values of these delayed rewards for each participant, and then subtracted 15 (value of the immediate reward) from the subjective value of each delayed reward, which resulted in the value difference between the immediate reward and the delayed reward (i.e., the no-brainer option). Using these value differences, we computed the predicted decision times for each participant, and we found that actual decision times did not significantly differ from predicted decision times (b = −0.01, SE = 0.008, t(264) = −1.86, p = .064, r = .11), suggesting that participants might be experiencing the predicted or expected amount of subjective conflict. That is, participants might have relied on subjective value computations to decide during no-brainer trials.

However, the analysis above provides only limited insights into whether participants were in fact computing subjective values. To further explore this question, we tested whether decision times during no-brainer trials varied as a function of delay (10, 30, or 60 days), with the logic being that if participants had indeed computed the subjective values of the no-brainer options, decision times should be faster as the delay increased. For example, the subjective value of $15 in 60 days is less than $15 in 10 or 30 days for all participants, and thus, the value difference between the immediate reward of $15 today and $15 in 60 would be the greatest and thus decision conflict should be the least (vs. $15 in 10 or 30 days). However, the relationship between decision time and delay was not significant (b = 0, SE = 0, t(117) = 0.47, p = .638, r = .04; 10 days: mean = 839 ms, SD = 33 ms; 30 days: 881 ms, SD = 34 ms; 60 days: 851 ms, SD = 33 ms), suggesting that decision times were similar when the delayed reward of $15 was presented in 10, 30, or 60 days. Thus, contrary to the analysis above, this finding suggests that participants might not have computed the subjective values of no-brainer options. That is, this result provides indirect evidence that participants might be doing something different on no-brainer trials, which might explain the different neurophysiological responses associated with these trials.

## 4. Discussion

We recorded EEG signals over midfrontal scalp electrodes and pupil dilation responses while participants made intertemporal decisions. As with recent work on theta-pupil relationships during inhibitory control tasks (Dippel et al., 2017; Mückschel et al., 2017), we show that midfrontal theta power and pupil dilation track subjective conflict during value-guided decisions involving trade-offs between costs and benefits occurring at different times. Even though participants were simply expressing their preferences for different intertemporal rewards and had little or no prepotent responses to inhibit, our results suggest that subjective conflict had parametrically modulated midfrontal theta power and pupil responses. Our study therefore extends previous work on conflict processing during inhibitory control and reinforcement learning tasks (Frank et al., 2015; Nakao et al., 2010; Yeung et al., 2004).

Both midfrontal theta and pupil responses track subjective intertemporal preferences, or how similar the subjective values of the immediate and delayed rewards were. The curvilinear effects suggest that these signals increased in magnitude when the two rewards had the same subjective value (high choice conflict); when the subjective value of one reward became higher than the other (less choice conflict), the magnitude of both signals decreased.

Unexpectedly but perhaps also the most interesting finding is that midfrontal theta and pupil responses were also enhanced when decisions were surprisingly easy (e.g., $15 today or $15 in 10 days), suggesting that these signals might track more than just conflict between two choices or responses. Instead, and consistent with past theorizing (e.g., Alexander and Brown, 2011; Cavanagh and Frank, 2014; Cavanagh and Shackman, 2015), these neurophysiological responses may signal events (e.g., surprising or conflicting) requiring increased attention and adaptive control, regardless of their valence (HajiHosseini and Holroyd, 2013; Hauser et al., 2014; Jahn et al., 2014; Sallet et al., 2013; Talmi et al., 2013).

When faced with a no-brainer choice, participants might realize that unlike other choices, it was unnecessary to consider the relevant costs (delay) and benefits (reward magnitude). They could instead rely on heuristics (e.g., always take the immediate reward), and the switch from value-to heuristics-guided decision strategy may be driven by adaptive control processes (Cohen, 2016; Holroyd and Coles, 2002; Karlsson et al., 2012; Shenhav et al., 2013; Tervo et al., 2014). Further, enhanced midfrontal theta and pupil responses might reflect dynamic switching between brain networks (Cocchi et al., 2013; Uddin, 2015). For example, past work suggests that pupil responses correlate strongly with LC-NE system activity (Aston-Jones and Cohen, 2005b; Murphy et al., 2014; Rajkowski et al., 1994), and this neuromodulatory system may be involved in resetting networks and optimizing behavior (Bouret and Sara, 2005; Nieuwenhuis et al., 2005; Sara and Bouret, 2012; Urai et al., 2017; Usher et al., 1999; Warren et al., 2016). Therefore, consistent with these suggestions, our results are also in line with the view that midfrontal theta oscillations and norepinephrine help implement adaptive control and optimize behavioral responses (e.g., Verguts and Notebaert, 2009).

Crucially, our findings provide evidence of convergence across qualitatively different sorts of stimuli—that midfrontal theta is involved in not only cognitive control during inhibitory control tasks (e.g., Stroop, go/no-go), reinforcement learning tasks (e.g., Cavanagh et al., 2014; Frank et al., 2015), but also intertemporal decisions that require participants to only express their pre-existing subjective preferences. Although our EEG electrode array preclude us from localizing midfrontal theta sources, previous work suggests that midfrontal theta oscillations are generated in the ACC and surrounding mPFC (Asada et al., 1999; Debener et al., 2005; Töllner et al., 2017). Thus, future work should not only investigate whether common neural sources generate midfrontal theta dynamics in value-guided choice and inhibitory control tasks, but also use value-guided choice paradigms to gain further insights into the functional significance of midfrontal theta dynamics. The latter is especially important, given that we are the first to investigate midfrontal theta during intertemporal choice and there is no one-to-one mapping between neural oscillatory frequencies and cognitive function. Instead, the theta dynamics we have observed may reflect memory load, mental effort, or binding of widely distributed cortical assemblies (Sammer et al., 2007; Wascher et al., 2014; Zakrzewska and Brzezicka, 2014). Such studies will help elucidate how midfrontal theta dynamics underlie good everyday self-regulation and decision making (e.g., Ertl et al., 2013; Knyazev 2007). Finally, because our main research question was how midfrontal theta—a canonical feature of performance and conflict monitoring during cognitive control tasks—also tracks conflict and adaptive control during value-guided choice, we have not examined oscillatory dynamics at other frequency bands that have also been associated with reward processing and value-guided choice (HajiHosseini & Holroyd, 2015; Polania et al., 2014, 2015). For those interested in exploring other frequency bands, our dataset can be downloaded from osf.io/7m9c2.

As for changes in pupil size, some have described these as reflecting changes in neural gain mediated by LC-NE system activity (Aston-Jones and Cohen, 2005b; Eldar et al., 2013; Murphy et al., 2014). For example, changes in pupil responses correspond to changes in locus coeruleus firing rate (Joshi et al., 2016; Murphy et al., 2014; Varazzani et al., 2015), as well as norepinephrine concentrations (Phillips et al., 2000; Warren et al., 2016). Extending recent work (Chmielewski et al., 2017; Dippel et al., 2017; Mückschel et al., 2017), we found that midfrontal theta correlated with pupil dilation responses, suggesting that the mPFC and LC-NE might interact to resolve conflict and respond to surprising events even during choices based on personal preferences. However, given that we did not directly measure mPFC and locus coeruleus activity, such suggestions remain speculative.

Future research using pharmacological interventions will be necessary to show that changes in pupil dilation and midfrontal theta during value-guided choice are indeed mediated by changes in norepinephrine concentrations. Such studies will help rule out other contributors to pupil responses, such as the colliculi (super and inferior) and other neurotransmission systems (dopamine and acetylcholine) (Sara 2009; Wang and Munoz, 2015). Interestingly, previous studies have found changes in neurophysiological activity (e.g., error-related negativity) but not behavioral responses or choices after manipulating norepinephrine concentrations (Jepma et al., 2010; Riba et al., 2005), indicating that more work needs to be done to understand the specific interactions between the LC-NE system and adaptive control processes such as theta oscillatory dynamics during decision making (Verguts and Notebaert 2009).

Because the intertemporal choice paradigm has generally been used to study self-control conflicts (e.g., Berns et al., 2007), our results suggest that overlapping processes may subserve inhibition processes and value-guided decision making, as well as self-control (Berkman, 2017; Berkman et al., in press; Shenhav, in press). For example, changes in midfrontal theta oscillations and pupil responses when faced with self-control conflicts involving cost-benefit trade-offs might reflect recruitment of the underlying adaptive control systems. As such, understanding how these systems are recruited and interact may explain why certain people are more successful than others at self-regulation.

In summary, we used economic modeling to show that subjective conflict during intertemporal choice parametrically modulated midfrontal theta power and pupil dilation responses. Unexpectedly, these signals also increased when intertemporal decisions were surprisingly easy, suggesting that these signals may instead reflect the need to increase attention and adaptive control to resolve conflicting or surprising events. Our results suggest that resolving choice conflict and exerting self-control during everyday choices may depend on interactions between neural systems that generate midfrontal theta oscillations and pupil dilation responses. Finally, our approach highlights the benefits of integrating cognitive neuroscience and neuroeconomics, which can provide insights into how inhibitory and adaptive control processes underlie value-guided choice. Conversely, neuroeconomic approaches offer paradigms that can inform and constrain cognitive neuroscience theories.

## Footnotes

1 A central research question in our lab is the relationship between cognitive control and negative affect, and the latter could potentially be indexed by activity in the corrugator supercilii muscles. Thus, we measure EMG activity in this muscle group in most of our EEG experiments (e.g., Elkins-Brown et al., 2016, 2017).

2 We previously used regression-based procedures (cf. Gratton, Coles, & Donchin, 1983) to correct for eye-blinks and obtained similar time-frequency and ERP results.

3 Granger tests have been useful in disciplines such as econometrics, but Granger estimates from neuroscience research can be severely biased or of high variance (Stokes and Purdon, 2017), and we consider this analysis to be highly exploratory.

## Acknowledgements

The authors thank Flavia Freitas Melcop Cardozo, Zahra Khan, and Jeyasakthi Venugopal for help with data collection and documentation. We also thank the anonymous reviewers, whose comments greatly improved the manuscript.

